# Identification of genes supporting cold resistance of mammalian cells: lessons from a hibernator

**DOI:** 10.1101/2023.12.27.573489

**Authors:** Masamitsu Sone, Nonoka Mitsuhashi, Yuki Sugiura, Yuta Matsuoka, Rae Maeda, Akari Yamauchi, Ryoto Okahashi, Junpei Yamashita, Kanako Sone, Sachiyo Enju, Daisuke Anegawa, Yoshifumi Yamaguchi

## Abstract

Susceptibility of human cells to cold stress restricts the use of therapeutic hypothermia and long-term preservation of organs at low temperatures. In contrast, cells of mammalian hibernators possess remarkable cold resistance, but little is known about the molecular mechanisms underlying this phenomenon. In this study, we conducted a gain-of-function screening of genes that confer cold resistance to cold-vulnerable human cells using a cDNA library constructed from the Syrian hamster, a mammalian hibernator, and identified Gpx4 as a potent suppressor of cold-induced cell death. Additionally, genetic or pharmacological inhibition of Gpx4 in a hamster cell line under prolonged cold culture led to cell death, which resembles ferroptosis characterized by accumulation of lipid peroxide and ferrous iron dependency. Genetic disruption of other ferroptosis-suppressing pathways, namely biopterin synthesis and mitochondrial or plasma membrane CoQ reduction pathways, accelerated cold-induced cell death under Gpx4 dysfunction. Collectively, ferroptosis-suppressing pathways protect the cells of a mammalian hibernator from cold-induced cell death and the augmentation of these pathways renders cold resistance to cells of non-hibernators, including humans.

## Introduction

Prolonged and severe low body temperature (Tb) is fatal to the majority of mammals. An exception to this rule is mammalian hibernation. Hibernation is an adaptive strategy to survive food-scarce seasons by lowering metabolism, thereby exhibiting extremely low body temperature (Tb) and saving energy expenditure (1). During the hibernation period, small mammalian hibernators (hereafter simply termed as “hibernators”) repeat many cycles of deep torpor and periodic arousal; in deep torpor, they cease motion and largely reduce heat production and their core Tb for several days to a couple of weeks; in periodic arousal, they recover their Tb to euthermia and sustain normal activity for less than one day (2). During deep torpor, their Tb drops to below 10°C and becomes stable at approximately 1°C above ambient temperature (3). Such very low Tb causes severe cellular and organ injuries in non-hibernators, including humans, mice, and rats, but not in small hibernators (4). However, it remains largely unknown how hibernators avoid cellular and organ injuries induced by very low Tb.

We previously reported that a hibernator, the Syrian hamster, Mesocricetus auratus (hereafter referred to as hamsters), is resistant to cold-induced ferroptosis in a nutrient-dependent manner. Primary hepatocytes derived from hamsters fed a diet containing a high amount of α-Tocopherol (αT) exhibit cold resistance by storing high amounts of αT in the cells, whereas those from hamsters fed a low-αT diet do not (5). In addition, primary hepatocytes isolated from mice fed a high-αT diet fail to store high amounts of αT and do not exhibit cold resistance (5). Thus, there is a difference in the mechanisms that exert αT-dependent cold resistance between hibernators and non-hibernators. Likewise, there is a difference in intrinsic, nutrient-independent, and cellular cold resistance between hibernators and non-hibernators. It has long been known that hibernators’ cells can survive longer than those from non-hibernators in cold temperatures lower than 10°C, and this was recently confirmed in cell lines derived from some hibernator species (6–9). However, little is known about the molecular mechanisms that support the cell-intrinsic cold resistance of hibernators.

Prolonged cold stress induces cell death in cells derived from non-hibernators including humans. Such cold-induced cell death fulfills the hallmarks of ferroptosis (10). Ferroptosis is a type of regulated cell death that accompanies ferrous-ion-mediated accumulation of lipid peroxide radical (11, 12). Along with normal cellular activities, lipid radicals are continuously produced by the oxidation of polyunsaturated fatty acids in the plasma membrane and are further oxidized into lipid peroxide radicals by molecular oxygen (13). Lipid peroxide radicals attack adjacent lipids to produce lipid radicals and lipid peroxide, and radical-trapping antioxidants (RTA) such as αT and CoQH_2_ can prevent this reaction (14). Depending on the ferrous ion, accumulated lipid peroxide is converted into lipid peroxide radicals, which propagate in the cell membrane and finally induce necrotic cell death (13).

Glutathione peroxidase (Gpx) family proteins reduce a variety of reactive oxygen species (ROS) by utilizing the reductive power of glutathione. Among eight members of Gpx genes in vertebrates, Gpx4 has unique characteristics in that it functions as monomer while many of other Gpx family members do as tetramer (15) and it has a catalytic center with more “open” structure than other Gpx proteins (16). Because of these characteristics, Gpx4 can reduce a wide variety of ROS, especially lipid peroxide, and plays a pivotal role in suppress ferroptosis (17). Most ferroptosis-inducing drugs developed to date directly or indirectly inhibit Gpx4’s function (17, 18). Gpx4 is ubiquitously expressed in the cytosolic and mitochondrial forms owing to its alternative transcriptional start sites (15). Loss of *Gpx4* in mice results in embryonic lethality (E7.5) (19), indicating that the removal of lipid peroxide by Gpx4 is essential for normal development.

Recently, multiple pathways that prevent ferroptosis in parallel with Gpx4 have been discovered: (i) the plasma membrane coenzyme Q (CoQ) reduction pathway: FSP1/Aifm2 reduces CoQ to CoQH_2_ depending on NAD(P)H, thereby indirectly reducing lipid peroxide radicals at the plasma membrane (20, 21). (ii) Biopterin pathway: Biopterin, a redox cofactor of several enzymes, can suppress the propagation of lipid peroxide radicals directly either or via αT (22). The expression levels of GCH1, the rate-limiting enzyme of the biopterin synthesis pathway, in human cancer cell lines are well correlated with resistance to ferroptosis-inducing drugs (23). (iii) Mitochondrial CoQ reduction pathway: DHODH is a mitochondrial enzyme that functions in the uridine synthesis pathway and indirectly reduces CoQ to CoQH_2_ to suppress the peroxidation of mitochondrial membrane (24). However, the functional contribution of these pathways, as well as Gpx4, to the intrinsic cold resistance of hibernators has not been investigated.

To elucidate the mechanisms of hibernators’ intrinsic cold resistance and to uncover potential approaches to bolster cold resistance in non-hibernators, including humans, we described metabolic changes during cold exposure and conducted a gain-of-function screening, in which cDNA derived from a cold-resistant hamster cell line was overexpressed in a cold-sensitive human cancer cell line to identify genes that confer cold resistance to human cells. Using a genome-editing approach and functional assays, we further examined the contribution of ferroptosis-suppressing pathways to cold resistance in mammalian cells.

## Materials and Methods

### Animals and housing

All animal care and experimental procedures were approved by the Ethics Committees of Hokkaido University (Ethical Approval no. 18-0140), and were conducted according to the ethics guidelines of Hokkaido University.

4-week old male Syrian hamsters were purchased from the breeder (SLC, Inc., Japan), fed an MR standard diet (Nihon Nosan, Japan), housed under summer-like conditions (light: dark = 14:10 h and ambient temperature = 22-25°C). For hibernation induction, approximately 10-week old male hamsters were transferred to and housed in a cold room under winter-like conditions (light: dark = 8:16 h and ambient temperature = 4-5°C). Typically, hamsters start hibernation within 3 months in a cold room. When the animals were immobile and had low Tb, they were judged to have deep torpor (DT). When the animals become spontaneously active after DT, they were judged as in periodic arousal (PA). 8-week old female BL/6N mice were purchased from the breeder (SLC, Inc) and fed an MR stock diet (Nihon Nosan) and housed under summer-like conditions. Animals were sacrificed by reperfusion under anesthesia with 4% isoflurane for hepatocyte isolation, or decapitation under anesthesia for liver sampling.

### Chemicals

The reagents used in this study were as follows: deferoxamine (Sigma, D9533), Ferrostatin-1 (Sigma, SML0583), idebenone (Tokyo Chemical Industry, C0134), mitoquinol (Cayman Chemical, 89950), BAPTA-AM (Tokyo Chemical Industry, T2845), RSL3 (Sigma, SML2234), BODIPY C11 (Thermo Fisher Scientific, D3861), ML210 (Cayman Chemical, 23282), BH2 (Cayman Chemical, 81882), methotrexate (Fujifilm Wako, 139-13571), and αTocopherol (Tokyo Chemical Industry, T2309).

### Cell culture

HapT1, HT1080, and HEK293T cells were cultured in DMEM supplemented with high glucose (Fujifilm Wako, 048-29763), 1× penicillin-streptomycin (Fujifilm Wako), 1×NEAA (Thermo Fisher Scientific), and 10% FBS (Nichirei, Japan). For cold culture, cells were seeded into 24-well plates (Corning or Iwaki) at 0.5 – 1 × 10^5^/well and incubated at 37 °C overnight. On the next day, the cells were transferred to and kept in a refrigerator set at 4 °C with culture medium supplemented with 0.1M Hepes pH7.4 (Thermo Fisher Scientific).

### Cell death assay

The amount of LDH in the supernatant was measured using the Cytotoxicity LDH Assay Kit-WST (Dojindo, Japan) following the manufacturer’s instructions. The activity of LDH in each sample was normalized to that of cells completely lysed with 1% Triton X-100. The absorbance was measured at 490 nm using Cytation5 (Agilent) after a 30-min incubation at 37 °C and adding Stop solution.

### Plasmids

To generate PB-TAC-BFP, we first generated PB-TAC-ERN-BFP by Gateway recombination between pENTR-tagBFP and PB-TAC-ERN (a gift from Knut Woltjen, Addgene #80475), after which the rtTA/NeoR expression cassette was deleted by BglII/NheI digestion, end blunting, and ligation. pT2K-EF1-rtTA-Neo was prepared using the In-Fusion HD cloning kit (Takara Bio) to recombine the rtTA/NeoR expression cassette excised by NotI/NheI from the PB-TAC-ERN and Tol2 backbone linearized by PCR, using pT2K-CAGGSv3-EGFP (25) as a template. To generate transfer plasmids of lentivirus vector for overexpression, coding sequences (CDS) of hamster Dhodh or Gch1 or N-terminally 3xFlag-taggged CDS of hamster FSP1 or sequence from the first methionine to poly-adenylation signal of cytoplasmic hs-or ma-Gpx4 were amplified using cDNA from HapT1 or HT1080 and inserted into PL-sin-EF1a-EiP (a gift from James Ellis) (26) by replacing EGFP. To generate plasmids to express Cas9 and sgRNAs to knockout hamster genes, guide RNAs were designed using CRISPRdirect (https://crispr.dbcls.jp/), and oligonucleotides containing the guide RNA sequences were inserted into the BbsI site of pX330 or BsmBI site of lentiCRISPR v2 (both gifts from Feng Zhang, Addgene#42230 and #52961, respectively)(27). To generate transfer plasmids of AAV for overexpression under the liver-specific TBG promoter, CDS of EGFP or a sequence from the first methionine to poly-adenylation signal of cytoplasmic maGpx4 were inserted into pENN.AAV.TBG.N-FLAG-mSTAT5bCA. WPRE.bGH (a gift from Rhonda Kineman, Addgene #184463) by replacing the N-FLAG-mSTAT5bCA.

### Expression library construction

The full-length cDNA expression library was constructed using the SMARTer RACE 5’/3’ Kit (Takara Bio, 634858) according to the manufacturer’s instructions for the In-Fusion SMARTer Directional cDNA Library Construction Kit (Takara Bio, 634933). 1µg of total RNA extracted from HapT1 with RNeasy Mini Kit (Qiagen) was annealed with 24pmol 3’ In-Fusion SMARTer CDS Primer (5’ - CGGGGTACGATGAGACACCATTTTTTTTTTTTTTTTTTTTVN–3’ where N = A, C, G, or T; V = A, G, or C) in 11µL, at 72 °C, 3min; 42 °C, 2min; 25 °C, 5min, followed by first strand synthesis with template switching by adding 9µL First-strand mix (4µL 5xFirst-Strand Buffer, 0.5µL 100mM DTT, 1µL 20mM dNTP, 1µL SMARTer II Oligonucleotide, 0.5µL Rnase Inhibitor, 2µL SMARTScribe Reverse Transcriptase) at 42 °C, 90min; 68 °C, 10min. cDNA was amplified in two tubes containing 50µL PCR reaction (1µL of first-strand cDNA, 25µL of PrimeSTAR MAX, 24pmol of 5’RACE Universal single primer, 24pmol of 3’ In-Fusion SMARTer PCR Primer; 5’ - CGGGGTACGATGAGACACCA-3, by 18 cycles of 98 °C, 10sec; 55 °C, 5sec; 72 °C, 90sec) followed by purification with MagExtractor (Toyobo). linearized PB-TAC plasmid was amplified in 50µL PCR reaction (5ng PB-TAC-BFP, 25µL of PrimeSTAR MAX, 10pmol of PB-TAC_cDNA_lib_IF_F2 primer; 5’-TGATACCACTGCTTGTTTGTACAAACTTGTGATGGCCG-3’, 10pmol of PB-TAC-ERN_cDNA_lib_IF_R primer; 5’-TCTCATCGTACCCCGAGCTAAAACGCGGCCTCGAATC-3,’ by 25 cycles of 98 °C 10sec; 63 °C, 5sec; 72 °C, 40sec, followed by purification with MagExtractor. cDNA was inserted into PB-TAC plasmid in 10µL of In-Fusion reaction (100ng of PCR-amplified cDNA, 100ng of linearized PB-TAC, 2µL of 5X In-Fusion HD premix) at 50 °C, 17min, followed by ethanol precipitation. NEB 10-beta Electrocompetent E. coli (New England Biolabs) was transformed with the resultant DNA of the In-Fusion reaction by GenePulserII (BioRad) at 2.0 kV, 200 Ω, 25µF, seeded on agar plates at 5,000 colonies/dish, and harvested the next day. The PB-TAC plasmid library was extracted using the Plasmid Midi Kit (Qiagen).

### Expression screening of the genes rendering cold resistance

The rtTA-expressing HT1080 cell line (HT1080 T2-ERN) was established by transfection of pT2K-EF1-rtTA-Neo and pCAGGS-T2TP, followed by G418 selection. HT1080 T2-ERN cells were seeded on two 100 mm dishes at 1 × 10^6^ cells/dish, and the next day, which is hereafter set as day0, transfected with a complex of 12µg PB-TAC plasmid library, 3µg pCX-IFP2 a PBase-expression vector (28), 18µL Lipfectamine3000 and 18µL P3000 reagent (Thermo Fisher Scientific) per dish. On day1, the cells were trypsinized and seeded in 20 dishes. On day2, 1µg/mL of Doxycycline (Sigma) was added until the end of the experiment. On day3, the cells were incubated in a refrigerator set at °C with culture medium supplemented with 0.1M Hepes pH7.4 (Thermo Fisher Scientific). On day9, the cells were incubated at 37°C, 5% CO2 in normal culture medium until they grew to semi-confluent cell density. The cells were treated with total three cycles of 6-day cold culture and 37 °C incubation, and lysed with 50µg/mL Proteinase K at 56 °C, 30min with shaking. Genomic DNA was extracted by phenol-chloroform extraction and isopropanol precipitation. Hamster genes inserted into the genome of surviving HT1080 were amplified in 50µL PCR reaction (1ng of genomic DNA, 25µL of PrimeSTAR MAX, 10pmol of 5’RACE Universal single primer (5’ - CAAGCAGTGGTATCAACGCAGAGT-3’), 10pmol of PB-TAC-ERN_cDNA_lib_IF_R primer) by 30 cycles of 98 °C, 10sec; 63 °C, 5sec; 72 °C, 90sec. The resultant DNA was separated by agarose gel electrophoresis, resulting in several bands of DNA fragments. These DNA bands were individually excised and purified with MagExtractor and sequenced with a 5’RACE Universal single primer using Sanger sequencing. To evaluate the ability of each candidate gene to render cold resistance, these DNA fragments were individually inserted into PB-TAC plasmid by In-Fusion reaction, and resultant plasmid was transfected into HT1080 T2-ERN with Lipofectamine3000 and bulk cells were used for cell death assay at 4 °C, under 1µg/mL of Doxycycline.

### VIKING method

Genetic disruption of Gpx4 by VIKING was performed as previous report (29). The principle of this method is to use drug resistance for selecting knockout cells in which a doner vector containing drug resistance gene is inserted into the target genomic loci cleaved by the gene-specific sgRNA and Cas9. In this process, the doner vector is linearized within the transfected cells by Cas9/sgRNA complex derived from a donor cleaving vector. Briefly, HapT1 cells were transfected with the donor vector (pVKG-Puro), donor cleaving vector (VKG1-gRNA-pX330), and pX330 vector expressing Gpx4-targeting sgRNA (sgGpx4) using PEI MAX (Polysciences). Transfected cells were sparsely reseeded and incubated with 2µg/mL puromycin (Sigma-Aldrich) for a week. The resulting colonies were manually picked and expanded without puromycin. Genetic disruption of Gpx4 was confirmed by genomic PCR using a primer set flanking the targeting site of sgGpx4.

### Immunoblot

Frozen liver tissues that were crushed using a Multi-beads Shocker (Yasui Kikai, Japan) or cultured cells were dissolved in RIPA buffer containing 1× cOmplete proteinase Inhibitor Cocktail (Roche). Protein concentration in the lysates was determined using a BCA Protein Assay Kit (Thermo Fisher Scientific). Protein samples (5 to 25µg/lane) were separated by 12.5% – 15% SDS-PAGE and transferred onto Immobilon-P PVDF membranes (Merck Millipore). Proteins were probed in Can Get Signal Solution1 (Toyobo) using following primary antibodies: anti-Gpx4 (1:10,000, Abcam, ab125066), anti-β-actin (1:20,000, CST, 4970S), anti-FSP1 (1:10,000, Proteintech, 20886-1-AP), anti-Dhodh (1:10,000, Proteintech, 14877-1-AP), anti-Gch1(1:10,000, Proteintech, 28501-1-AP), anti-Spr (1:10,000, Proteintech, 16822-1-AP). HRP-conjugated anti-rabbit-IgG antibody (1:20,000, Jackson ImmunoResearch, 111-035-003) in Can Get Signal Solution2 (Toyobo), Immobilon Western Chemiluminescent HRP Substrate (Merck Millipore), and ImageQuant LAS4000 (GE Healthcare) were used for detection.

### BODIPY C11 assay

HT1080 and HapT1 cells were seeded at a density of 2 × 10^4^ cells/well in 96 well plates. next day, medium was replaced with culture medium containing 10µM BODIPY C11 (Thermo Fisher Scientific), and cells were incubated in 5% CO2 at 37 °C for one hour. The cells were washed three times with prewarmed HBSS(+) and the medium was replaced with HBSS(+) supplemented with 0.1M HEPES pH7.4 with or without 2µM RSL3 or 1µM Ferrostatin-1. Then, the cells were incubated in a refrigerator at 4 °C and fluorescent signal of BODIPY C11 was measured everyday with ex/em = 580/590 for reduced form and ex/em = 488/510 for oxidized form a Cytation5 plate reader in a monochromator mode. Time-lapse images was taken at 4 °C in a cold room using BZ-X800 (Keyence) fluorescent microscope.

### Sample preparation for metabolomic and lipidomic analysis

WT or Gpx4 KO HapT1 cells were seeded onto 100mm dishes at 3×10^6^ cells/dish. On the next day, cells for day0 samples were washed with HBSS(+) supplemented with 0.1M Hepes pH7.4, collected with cell scraper, centrifuged and resulting pellets were stored at −80°C until further use. The remaining cells were transferred to and kept in a refrigerator set at 4 °C with HBSS(+)/0.1M Hepes and processed on the corresponding day as same with day0 samples.

### Metabolomic Analysis

Methanol (500 μL) containing internal standards (IS: Methionine sulfone and 2-morpholinoethanesulfonic acid were used as IS for cationic and anionic metabolites, respectively) was added to the frozen cell pellets, followed by sonication and the addition of half volume of ultrapure water (LC/MS grade, Wako Pure Chemicals, Tokyo, Japan) and 0.4 volume of chloroform. The suspension was centrifuged at 15,000 x g for 15 min at 4°C. The resulting upper (aqueous) and lower (organic) layers were used for hydrophilic metabolome and lipidome analysis, respectively. For hydrophilic metabolome analysis, the aqueous layer was filtered through ultrafiltration tubes (Ultrafree MC-PLHCC, Human Metabolome Technologies, Tsuruoka, Japan), and the filtrate was dried up under nitrogen gas flow. The concentrated filtrate was dissolved in 70 μL ultrapure water. Anionic and cationic metabolites were quantified by IC-MS and LC-MS/MS, respectively.

Anionic metabolites, including sugar phosphates, organic acids, and nucleotides, were analyzed by IC-MS. An orbitrap MS system (Q-Exactive focus, Thermo Fisher Scientific) connected to a high-performance IC system (ICS-5000+, Thermo Fisher Scientific) was used for metabolite detection. The IC system was equipped with an anion electrolytic suppressor (Dionex AERS 500; Thermo Fisher Scientific) to convert the potassium hydroxide gradient to pure water before the sample entered the MS system. Separations were performed using a Thermo Fisher Scientific Dionex IonPac AS11-HC 4 μm particle size column with an IC flow rate of 0.25 mL/min, supplemented with a 0.18 mL/min post-column MeOH makeup flow. Potassium gradient conditions were as follows: 1 mM to 100 mM (0-40 min), 100 mM (40-50 min), and 1 mM (50.1-60 min) at a column temperature of 30 °C. The Q-Exactive focus mass spectrometer was operated in ESI-positive-negative switching mode. Full mass scans (m/z 70-900) were performed at 70,000 resolution. The automatic gain control target was set to 3 × 10^6 ions with a maximum ion injection time of 100 ms. The ionization parameters of the ion source were optimized: spray voltage of 3 kV, transfer tube temperature 320 °C, S-lens level 50, heater temperature 300 °C, sheath gas 36, auxiliary gas 10.

Cationic metabolites, including amino acids and nucleosides, were quantified using a triple quadrupole mass spectrometer (LCMS-8060, Shimadzu) equipped with an electrospray ionization (ESI) ion source in positive and negative ESI, multiple reaction monitoring (MRM) mode. Samples were separated on a Discovery HS F5-3 column (2.1 mm i.d. × 150 mm, particle size 3 μm, Sigma-Aldrich) using mobile phase A (0.1% formic acid/water) and mobile phase B (0. 1% formic acid/acetonitrile) in the ratios 100:0 (0-5 min), 75:25 (5-11 min), 65:35 (11-15 min), 5:95 (15-20 min) and 100:0 (20-25 min) with a stepwise gradient at a flow rate of 0.25 ml min-1 and a column temperature of 40 °C. Chromatogram peaks obtained in compound specific MRM channels were integrated and manually confirmed. Peak quantitation values obtained for each compound were corrected for recovery by IS and cell number.

### Detection of lipid peroxidation by LC-MS

400 u L of the lower layer (organic layer) was aliquoted into a separate tube, completely dried with nitrogen gas, redissolved in 70 uL MeOH and measured by LC-MS. For lipidomic analysis, an Orbitrap MS (Q-Exactive Focus, Thermo Fisher Scientific, San Jose, CA) connected to HPLC (Ultimate 3000 system, Thermo Fisher Scientific) was used. LC and MS conditions were as described by Růžička et al (30). Briefly, samples were injected into a Thermo Scientific Accucore C18 column (2.1 × 150 mm, 2.6 μm); mobile phase A was 10 mM ammonium formate, 50% acetonitrile (v), 0.1% formic acid (v); mobile phase B was acetonitrile/2 mM ammonium formate and isopropyl alcohol/water, 10:88:2 (v/v/v), 0.02% formic acid (v). The gradient elution profiles were as follows: 65:35 (0 min), 40:60 (0-4 min), 15:85 (4-12 min), 0:100 (12-21 min), 0:100 (21-24 min), 65:35 (24-24.1 min), 100:0 (24.1-28 min). The Q-Exactive Focus mass spectrometer was operated in both ESI positive and negative modes. After a full mass scan (m/z 250-1100), three data-dependent MS/MS runs were performed at 70,000 and 17,500 resolution, respectively. The automatic gain control target was set to 1 × 10^6 ions and the maximum ion injected time was 100 ms. Source ionization parameters were as follows: spray voltage 3 kV, transfer tube temperature 285 °C, S-lens level 45, heater temperature 370 °C, sheath gas 60, auxiliary gas 20. LipidSearch software (Mitsui Information, Tokyo, Japan) with the following parameters: precursor volume tolerance = 3 ppm, product volume tolerance = 7 ppm, m-score threshold = 3.

### Lentivirus vector production and infection

HEK293T cells (4.5 × 10^6^) were seeded onto a 100 mm dish, and the next day, were transfected with a complex of 5µg of pCMV-VSV-G (RIKEN BRC, RDB04392), 5µg of psPAX2 (a gift from Didier Trono, Addgene #12260), 5µg of transfer plasmid and 100µg of PEI MAX. Three days after transfection, culture supernatant was recovered and mixed with a third volume of LentiX Concentrator (Takara Bio), incubated at 4°C overnight, and centrifuged at 1,500g for 45min. The resulting pellet was dissolved in PBS and stored at −80°C. For infection, cells were seeded on plates, and the next day, they were incubated with lentivirus in the presence of 8µg/mL polybrene (Nacalai Tesque) for one day. The next day, the medium was changed and supplemented with 1µg/mL (HT1080) or 2µg/mL (HapT1) of puromycin. After 2-to 3-day treatment, cells were reseeded and cultured without puromycin.

### AAV vector production and infection

AAV was prepared according to a previous report with some modifications(31). HEK293T cells were seeded onto 100 mm dishes at 4.5 × 10^6^ /dish, and the next day, were transfected with a complex of 4µg of pAAV2/8 (a gift from James M. Wilson, Addgene #112864), 4µg of pHelper (Takara Bio), 4µg of transfer plasmid and 48µg of PEI MAX per dish. The next day, the medium was changed to DMEM without serum, and the cells were incubated for 5 days. The culture supernatant was recovered, centrifuged to remove cell debris, supplemented with a fourth volume of 40% polyethylene glycol (MW8000, Sigma)/2.5M NaCl, and incubated on ice for more than two hours. virus was precipitated by centrifugation at 3200×g, 4 °C for 30min, dissolved in 1mL/dish PBS containing 2.5mM MgCl_2_, and treated with 0.25µL/mL Benzonase (Sigma, E1014) at 37 °C, 30min. The reaction was stopped by adding 1mM EDTA. Then, saturated (NH_4_)_2_SO_4_ per 1mL of the virus solution and centrifuged at 9000 g, RT for 5 min to remove debris. The virus solution was filtered through a 0.45µm NEW Steradisc25 (Kurabo, Japan) and subjected to ultrafiltration with Amicon Ultra-15, 100kDa (Merck Millipore) by centrifugation at 3200×g, RT, followed by three PBS washes. filtered and concentrated virus solution was centrifuged at 15000 rpm for 5min to remove debris and stored at −80 °C. The virus titer was quantified using the AAVpro Titration Kit (for Real Time PCR) Ver.2 (Takara Bio) according to the manufacturer’s instructions.

For infection, 8 to 12-week old C57BL/6N female mice were intraperitoneally injected with 2.5 × 10^12^vg AAV. Four weeks after infection, mice were subjected to primary hepatocyte preparation.

### Primary culture of hepatocytes

Murine hepatocytes were isolated and cultured as previously described(5). Briefly, under anesthesia with 4% isoflurane, the livers were perfused from the portal vein with a solution containing 1mM EGTA in Ca^2+^/Mg^2+^-free HBSS, followed by a solution containing 1mg/mL collagenase and Ca^2+^/Mg^2+^ in HBSS at 37°C. Hepatocytes were mechanically dissociated in EMEM (Sigma Aldrich, M4655) containing 10% FBS, filtered through a 100µm cell strainer, and collected by centrifugation at 40 × g for 1 min. The cells were then resuspended in medium containing Percoll (GE Healthcare), centrifuged at 60 × g for 10 min, washed twice in EMEM containing 10% FBS, and finally filtered with a 40μm cell strainer.

The basal medium for culturing hepatocytes was DMEM/F12 (Gibco 21041-025) supplemented with 5mM HEPES pH7.4, 30 mg/L L-proline, 5 g/L BSA (Sigma A1470), 10ng/mL EGF (Sigma E4127), 1× Insulin, Transferrin, Selenium, Ethanolamine Solution (ITS-X) (Thermo Fisher Scientific), 0.1mM dexamethasone (Fujifilm Wako), 10mM nicotinamide (Fujifilm Wako), 1mM L-ascorbic acid 2-phosphate (Fujifilm Wako), and 1 × Penicillin-Streptomycin (Fujifilm). Hepatocytes were seeded in 24-well plates coated with atelocollagen (KOKEN, IPC-30) in basal medium supplemented with 10% FBS at 5 × 10^4^ cells/well and cultured at 37 °C. Three to four hours after seeding, the medium was replaced with serum-free basal medium. The next day, medium was replaced with basal medium supplemented with DMSO or 20µM BH2 or 10µM αTocopherol (αT) for 1hr at 37 °C. The medium was replaced with the basal medium containing 100mM HEPES pH 7.4 and DMSO/BH2/αT, and the cells were incubated at 4 °C in refrigerator.

### qRT-PCR

cDNA was synthesized from the total RNA of HapT1 cells using the PrimeScript RT Reagent Kit (Takara Bio). The amounts of Pts and Actb were quantified on a LightCycler 480 (Roche) using TB Green Premix ExTaq II (Takara Bio) with the following primers: Pts-F(5’-TACATGGAGGAGGCCATCAT-3’), Pts-R(5’-TCTGTTGTGCTTACAACGTC-3’), Actb-F(5’-AAGGCCAACCGTGAAAAGAT-3’), and Actb-R(5’-CCAGAGGCATACAGGGACAG-3’).

## Results

### Sustained glutathione synthesis pathway in Syrian hamster cells under cold conditions

The Syrian hamster pancreatic cancer cell line HapT1 survived for 5 days under cold (4°C) and nutrient-deprived culture conditions. Metabolomic analysis was performed to gain insight into the metabolic responses of HapT1 cells under these conditions. Hydrophilic metabolome profiling revealed continuous changes during the 7-day cold exposure period as indicated by PLS-DA and hierarchical clustering analysis (Fig. 1a). Hydrophilic metabolites could be classified into three clusters based on their response to cold exposure: (cluster-1) metabolites that rapidly and drastically decreased after cold exposure; (cluster-2) metabolites that transiently increased after cold exposure and then gradually decreased; and (cluster-3) metabolites that increased after cold exposure and peaked on day 5 (Fig. 1b). Most of the amino acids were classified in cluster 1, indicating that amino acids were depleted on the first day of transfer to cold and nutrient-deprived culture conditions (Fig. 1c). In addition, there was a significant reduction in mitochondrial electron carriers, namely NAD and NADH, and a reducing equivalent, NADPH (Fig. 1d), implying a decrease in the energy metabolic pathways. In contrast, the intracellular levels of high-energy nucleotides, including ATP, were maintained until day 5 (Fig. 1e). Focusing on the glutathione synthesis, reduced glutathione (GSH) did not show a considerable decline until day 5, while oxidized glutathione (GSSG) displayed a notable rise by day 2 (Fig. 1f). The levels of intermediate metabolites involved in glutathione synthesis remained relatively stable (homocysteine and cystathionine) or increased (cysteine). Consistent with this, methionine, the amino acid source for cysteine and glutathione synthesis, was rapidly and gradually depleted after cold exposure (Fig. 1f). These data suggest that the continuous supply of substrates for glutathione synthesis may allow the glutathione redox system to withstand possible oxidative stress during cold exposure in HapT1 cells.

**Fig. 1.**
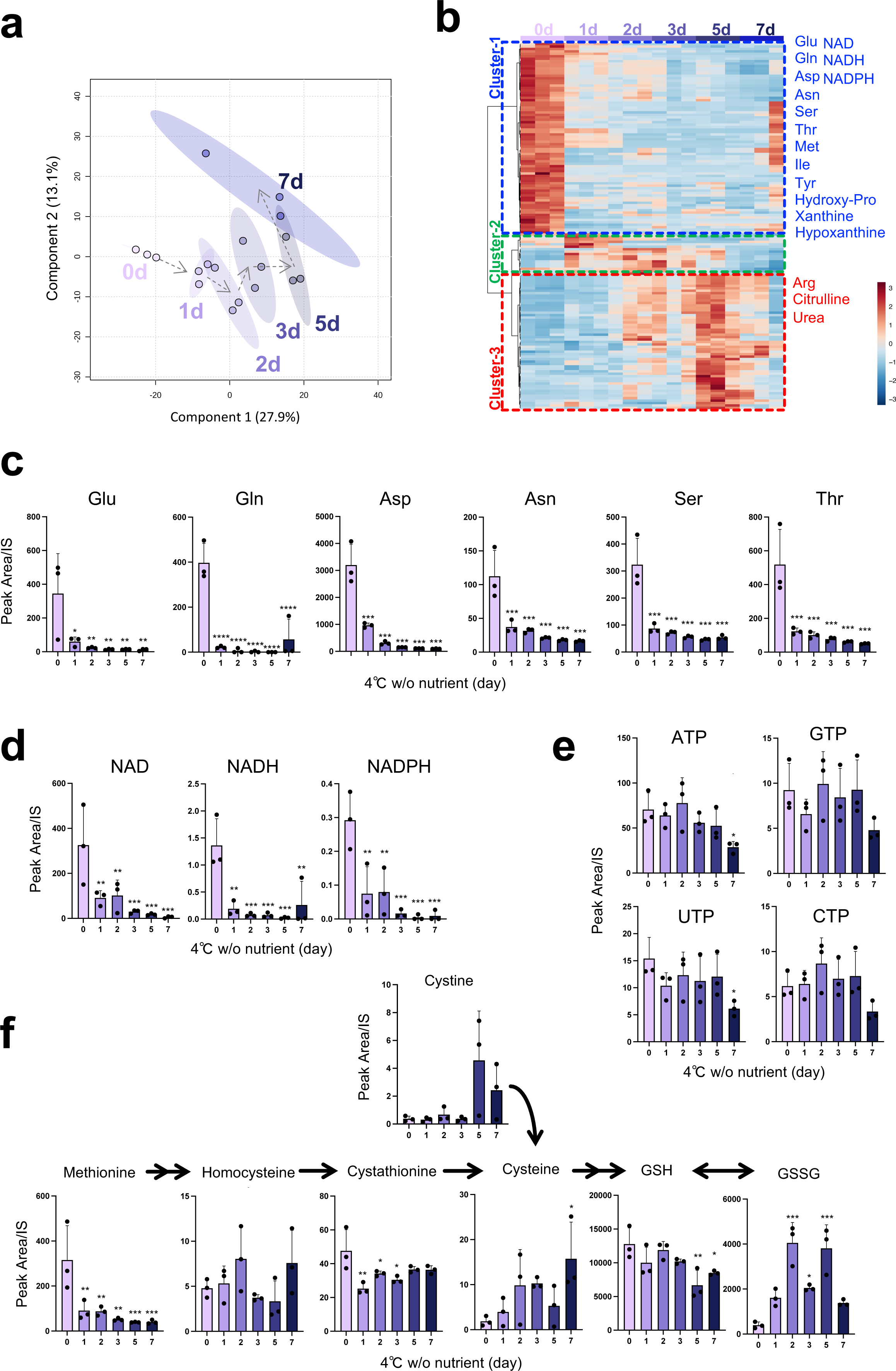
Metabolome analysis revealed glutathione synthesis pathway is retained in hamster cells under cold and nutrient-deprived condition. (A) PLS-DA analysis of metabolome profiles. 0d, 1d, 2d, 3d, 5d, and 7d indicate days (d) under cold temperature (4°C) and nutrient deprivation. Each dot corresponds to each sample (N=3 per day). (B) Hierarchical clustering of metabolome profiles. Representative metabolites are shown. (C) Amount of amino acids before or under cold temperature (4°C) and nutrient deprivation. (D) Amount of NAD, NADH, NADPH before or under cold temperature (4°C) and nutrient deprivation. (E) Amount of nucleotides before or under cold temperature (4°C) and nutrient deprivation. (F) Amount of metabolites related to glutathione synthesis before or under cold temperature (4°C) and nutrient deprivation. One-way ANOVA with the Dunnett’s multiple comparison test compared to day0 in C-F, *p < 0.05, **p < 0.01, ***p < 0.001, ****p < 0.0001

### Gain-of-function screening identified Gpx4 as a suppressor of cold-induced cell death in human cancer cell lines

In contrast to HapT1, the human fibroma cell line HT1080 died within 2 days of cold culture (Fig. 2a). We hypothesized that HapT1 expresses genes that can confer long-term cold resistance not only to hamster cells but also to cold-vulnerable human cells if exogenously introduced. To identify such genes, we conducted expression screening in which a full-length cDNA library prepared from HapT1 was stably introduced and expressed in a cold-vulnerable HT1080 via the piggyBAC transposon system (Fig. 2b). The expression of introduced genes was induced by doxycycline (Dox) under the control of the Tet-ON system. The bulk of the HT1080 cells were exposed to repeated cycles of 4°C for 6 days and 37°C for several days, which mimics the situation of deep torpor-arousal cycles. This treatment hardly allowed the survival of the cells without Dox but recovered the cell population in the presence of Dox, implying that the recovered cells acquired resistance to cold-rewarming stress by overexpression of hamster genes integrated into its genome. We then extracted genomic DNA from the surviving cells and analyzed the hamster genes that were stably integrated in the cells (Fig. 2b). As a result, 15 genes were identified, and only Gpx4 was inserted into the genome of surviving cells in three independent screening trials (Fig. 2c). In addition, when each of the 15 hamster genes was independently introduced and exogenously expressed in HT1080, only Gpx4 provided long-term cold-resistance to HT1080 (Fig. 2c, 2d).

**Fig. 2.**
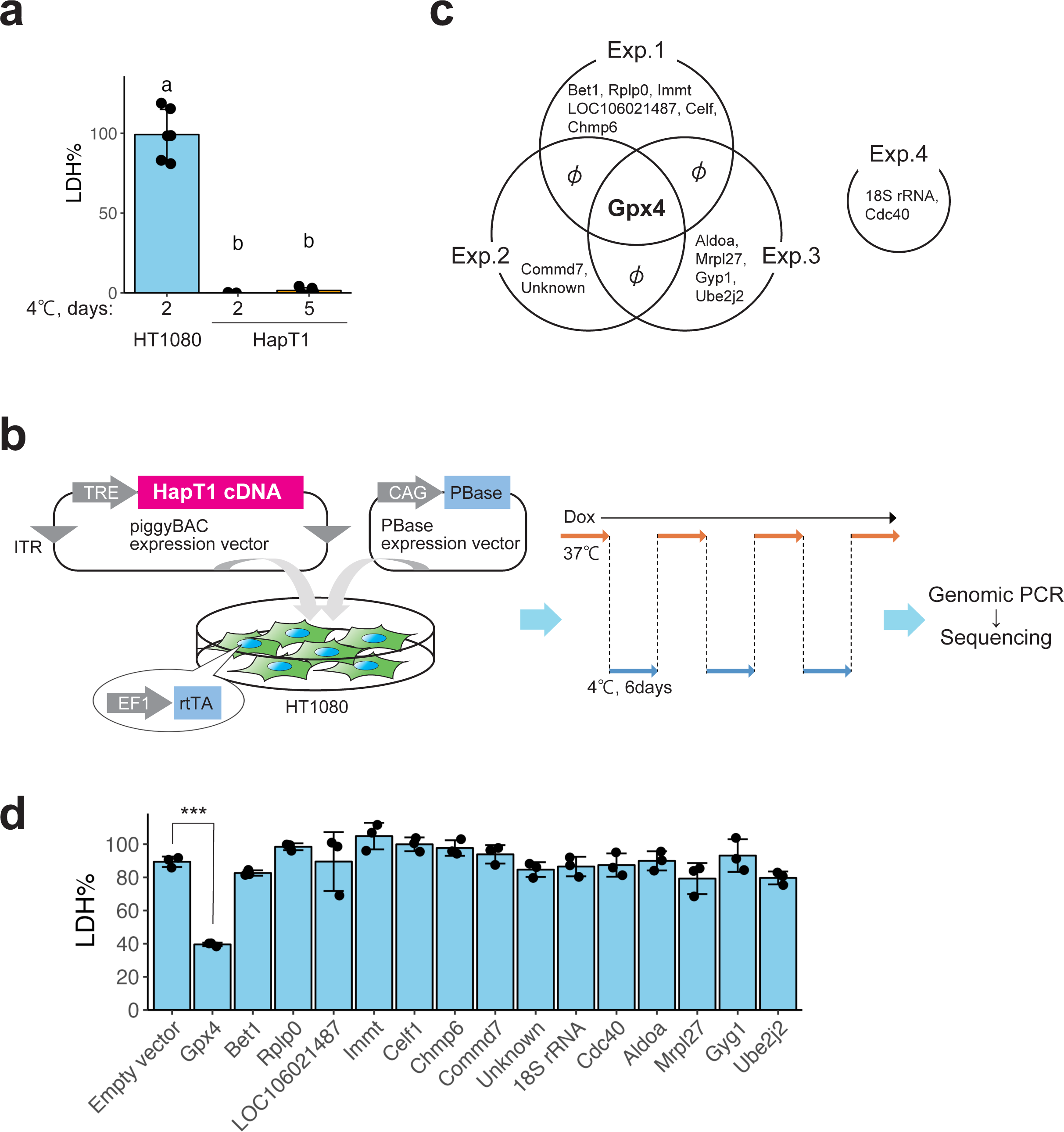
Gpx4 was identified by hamster cDNA expression screening as a gene rendering cold-resistance to a human cancer cell line. (A) Cold resistance of hamster HapT1 cells but not of human HT1080 cells. The proportion of dead cells was determined by LDH assay (One-way ANOVA with the Tukey’s multiple comparison test, p < 0.05). (B) Schematic illustration of hamster cDNA expression screening for genes that render the resistance to cold-rewarming culture to human HT1080 cells. (C) Venn diagram of the genes inserted in the genome of survived cells under cold-rewarming culture in four screening experiments. (D) The proportion of dead cells in HT1080 cells after 24hr of cold (4°C) culture when each indicated gene was overexpressed (N = 3 wells, One-way ANOVA with the Dunnett’s multiple comparison test, ***p < 0.001).

### Loss of Gpx4 function abolishes long-term cold resistance in hamster cells due to the accumulation of lipid peroxide

We next addressed whether Gpx4 was required for cold resistance in HapT1 cells. Three independent Gpx4 knockout (KO) HapT1 lines were established using the CRISPR-based VIKING method (29) (Fig. 3a). These Gpx4 KO lines proliferated without obvious abnormalities at 37°C (data not shown). When exposed to cold (4°C), most Gpx4 KO cells survived for 2 days, but died by day5 (Fig. 3b). Cell death in Gpx4 KO HapT1 cells induced by prolonged cold stress was prevented by ferroptosis inhibitors, iron chelator (deferoxamine), lipophilic antioxidant (Ferrostatin-1), and coenzyme Q mimetics (idebenone and mitoquinol) (Fig. 3c). In addition, similar to a previous report on human cells (32), a calcium chelator (BAPTA-AM) inhibited cell death of Gpx4 KO HapT1 cells in cold conditions (Fig. 3c, See Discussion). We then examined another hallmark of ferroptosis, lipid peroxidation, using live-imaging analysis with BODIPY C11. This analysis revealed that lipid peroxidation in cold culture was much slower in WT HapT1 cells than that in HT1080 cells (Fig. 3d, 3e). Gpx4 KO HapT1 cells exhibited a much faster accumulation of lipid peroxide than parental (wild-type; WT) HapT1 cells, whereas both Gpx4 KO and WT HapT1 cells exhibited very low levels of lipid peroxidation before cold culture (Fig. 3d, 3e). Extensive lipid peroxidation by cold culture was also observed in WT HapT1 cells treated with a direct Gpx4 inhibitor, RSL3, at a rate comparable to that in Gpx4 KO lines (Fig. 3e). Consistent with this, oxidized lipids, oxidized phosphatidylethanolamines (PE38:4;O, PE38:4;O2, and PE38:4;O3), which are ferroptosis signatures (33), increased in Gpx4 KO HapT1 cells after cold treatment (Fig. 3f). These results suggested that endogenous Gpx4 acts as a suppressor of cold-induced ferroptosis in HapT1 cells. To examine which period of cold culture Gpx4 exerts its function in preventing cold-induced ferroptosis, we treated WT HapT1 cells with RSL3 at different time windows (Fig. 3g) and quantified the amount of cell death. This experiment demonstrated that duration of Gpx4 dysfunction in cold was well correlated with the proportion of dead cells, and that Gpx4 inhibition at 37°C before cold culture did not affect the cold resistance of the cells. Therefore, the function of Gpx4 is required at 4°C, but not at 37°C, to avoid cold-induced cell death. Taken together, these results suggest that Gpx4 functions in cold culture to confer cellular resistance against cold-induced ferroptosis.

**Fig. 3.**
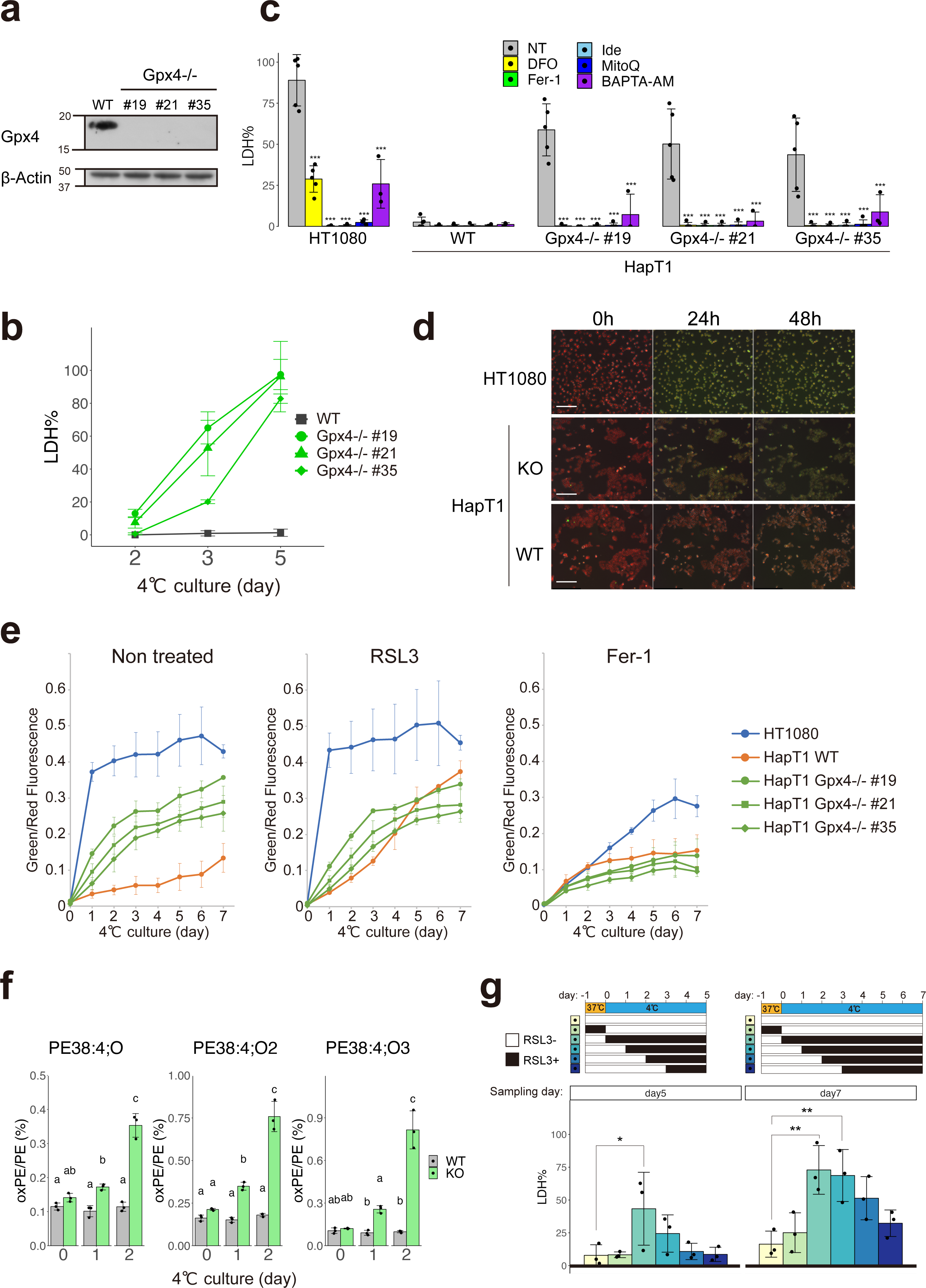
Gpx4 is necessary for preventing lipid peroxidation and cell death induced by long-term cold exposure in hamster HapT1 cells. (A) Immunoblot of Gpx4 and β-Actin proteins in parental (wild type; WT) and clonal HapT1 cells (#19, #21, #35) in which Gpx4 gene is disrupted. (B) The proportion of dead cells in WT or Gpx4-KO HapT1 cells during cold (4°C) culture. (C) The proportion of dead cells in HT1080 and parental or Gpx4-KO HapT1 cells after 5 days of cold (4°C) culture in the absence (non-treated; NT) or presence of 100µM DFO, 1µM ferrostatin-1 (Fer-1), 10µM idebenone (Ide), 10µM mitoquinol (MitoQ), 10µM BAPTA-AM (One-way ANOVA with the Dunnett’s multiple comparison test compared to NT within each cell lines, ***p < 0.001). (D) Oxidized lipid detection by BODIPY C11 ratio imaging in HT1080 and HapT1 cells (parental or Gpx4-KO) during cold (4°C) culture. Scale bar = 200µm. (E) Time-course changes of oxidized lipid level determined with BODIPY C11 ratio imaging in the presence or absence of 2µM RSL3 or 1µM Fer-1 N = 3. Plot is represented as mean±s.d. (F) Detection of lipid peroxidation by LC-MS/MS analysis. The amount of mono-(left), di-(middle), tri-(right) oxidized phosphatidylethanolamine (PE38:4) in WT or Gpx4-KO HapT1 cells are shown (One-way ANOVA with the Tukey’s multiple comparison test, p < 0.05). (G) The proportion of dead cells in HapT1 after 5 or 7 days of cold (4°C) culture in the absence or presence of 2µM RSL3. Each bar corresponds to the condition shown in the upper panel wherein RSL3 was added only during the period indicated by black during the culture (One-way ANOVA with the Dunnett’s multiple comparison test, *p < 0.05, **p < 0.01).

### Both hamster and human Gpx4 have potential to render cold resistance

The result that misexpression of hamster Gpx4 (maGpx4) increased the survival rate of HT1080 under cold conditions raises the possibility that maGpx4 is more effective in preventing cold-induced ferroptosis than human Gpx4 (hsGpx4), because HT1080 expresses comparable amounts of endogenous hsGpx4 (Fig. 2d, Fig. S1a). To test this possibility, we overexpressed either maGpx4 or hsGpx4 in HT1080 cells using lentivirus vectors and examined its effect on cold resistance (Fig. 4a). Overexpression of each Gpx4 gene provided cold resistance to human cells to the same extent (Fig. 4b). In addition, either of these Gpx4 genes restored long-term cold resistance in Gpx4 KO HapT1 cells (Fig. 4a, 4c). In summary, these results suggest that maGpx4 and hsGpx4 have comparable abilities to prevent cold-induced ferroptosis when overexpressed.

**Fig. 4.**
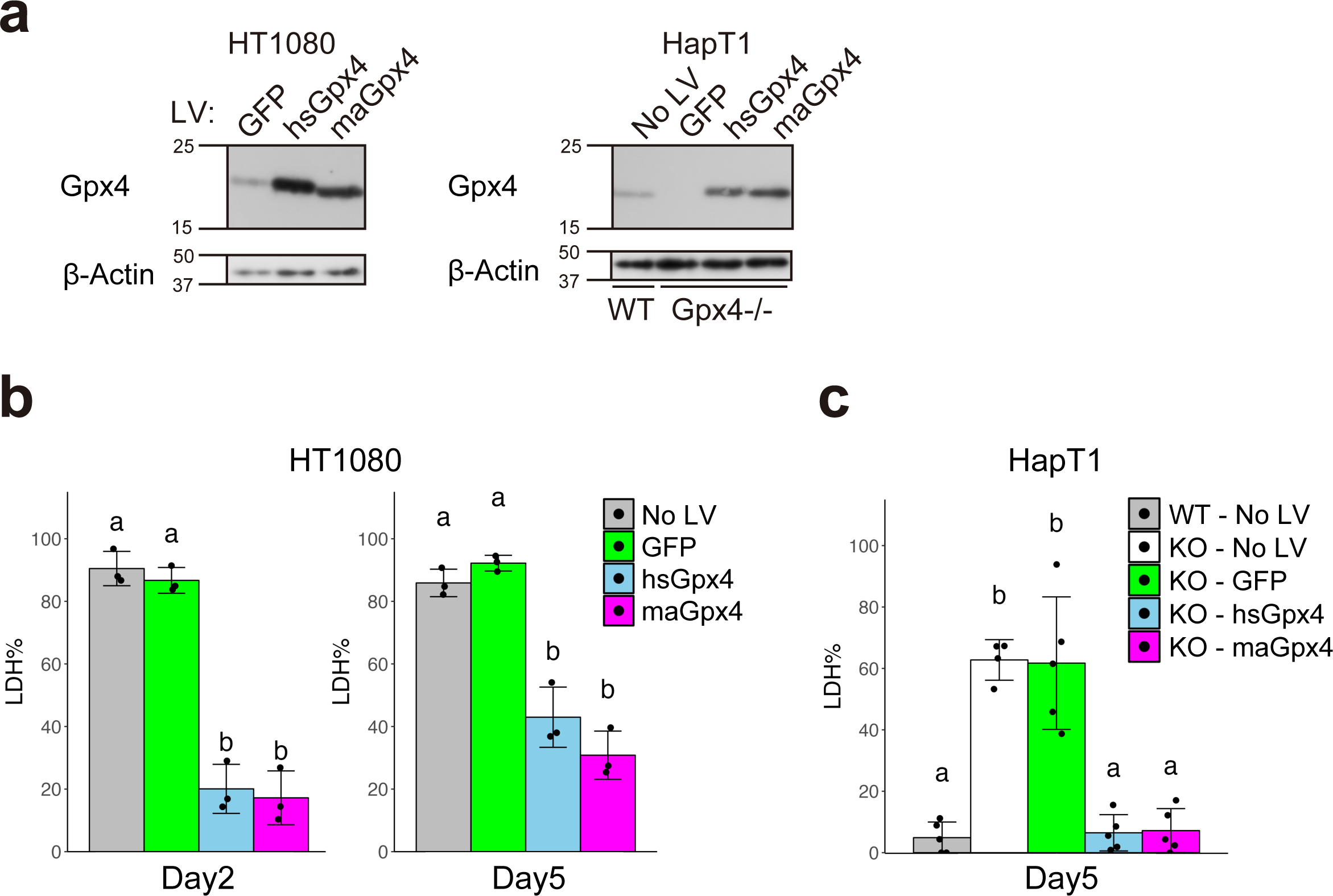
Overexpression of either human or hamster Gpx4 prevents cold-induced cell death. (A) Immunoblot of Gpx4 and β-Actin proteins in HT1080 and HapT1 (WT and Gpx4-KO clone #21), in which GFP, human (hs)Gpx4, or hamster (ma)Gpx4 were exogenously expressed by lentivirus vectors at 37°C. (B, C) The proportion of dead cells after 5 days of cold (4°C) culture in HT1080 (B) and HapT1(WT or Gpx4-KO) (C), in which lentivirus vectors expressing GFP or hs/maGpx4 were infected (One-way ANOVA with the Tukey’s multiple comparison test, p < 0.05).

Based on the finding that higher Gpx4 expression provides higher cold resistance in both hibernator and non-hibernator cells, we investigated the expression levels of Gpx4 in the livers of hamsters and mice. The amount of this protein was comparable between the two species (Fig. S1b). Thus, abundance of Gpx4 did not explain the cold-resistance difference between hamsters and mice. In addition, the expression levels of Gpx4 protein in the hamster liver did not change throughout the seasons (Fig. S1c), excluding the possibility that cold resistance is regulated by the regulation of Gpx4 protein level, at least in the liver, during hibernation.

### Biopterin and CoQ reduction pathways support cold resistance of mammalian cell lines parallel to Gpx4

Although Gpx4 KO HapT1 cells lost long-term cold resistance, they survived for at least two days under cold conditions and therefore can be considered to possess short-term cold resistance (Fig. 3b). To examine whether HapT1 cells possess redundant pathway(s) other than Gpx4 to prevent cold-induced lipid peroxidation and ferroptosis, we knocked out other known ferroptosis suppressors, namely, FSP1, Dhodh, and Gch1, using the lentiviral CRISPR/Cas9 system (Fig. 5a). A single disruption of each of these genes did not affect the survival rate of HapT1 cells under cold (Fig. 5b-d). However, when treated with Gpx4 inhibitors (RSL3 or ML210), these KO cells died at a significantly higher rate than WT cells by 2 or 3 days under cold (Fig. 5b-d, S2a-c). These data suggest that each of these genes contributes to cold resistance, in parallel with Gpx4.

**Fig. 5.**
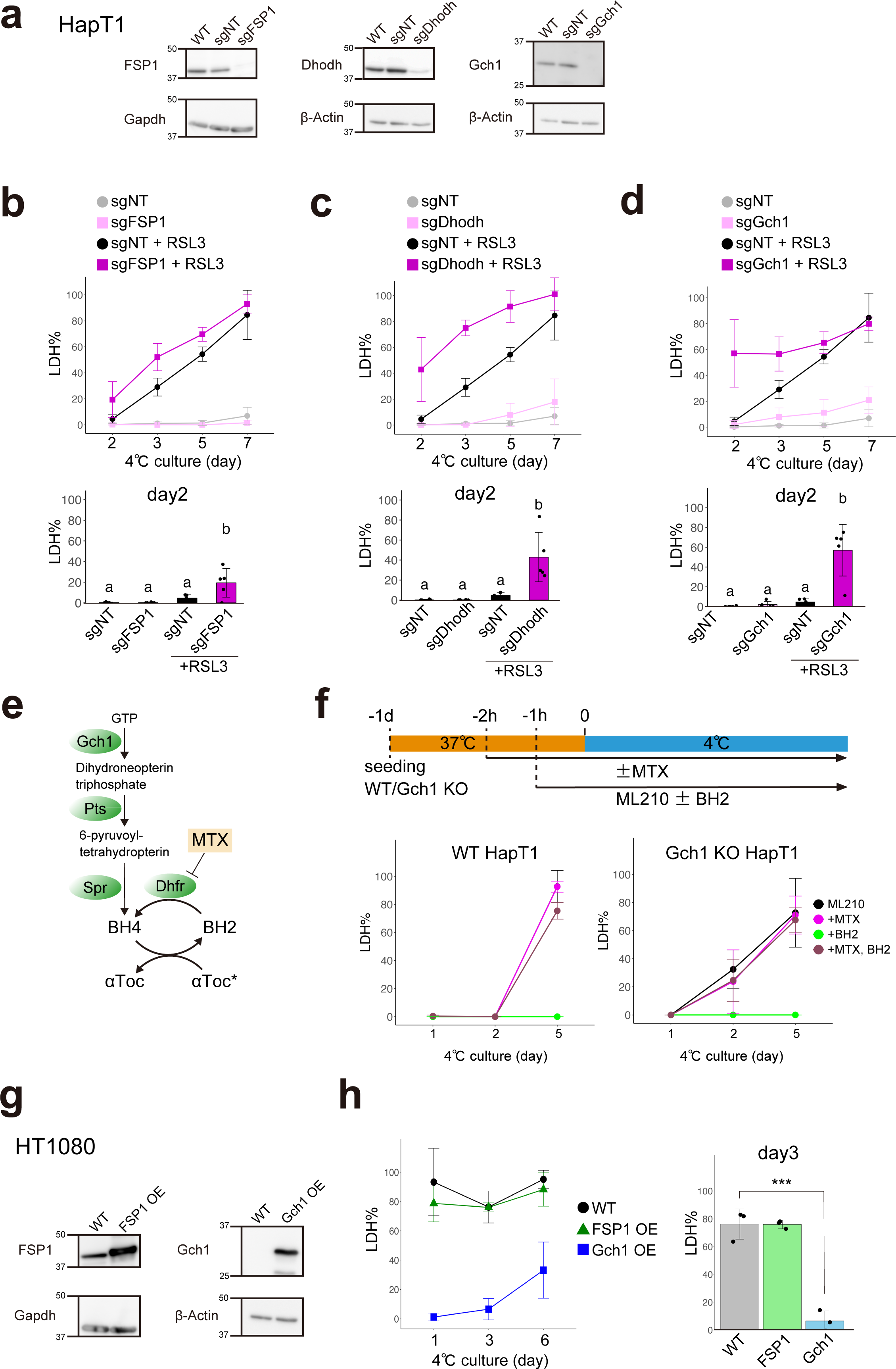
Biopterin and CoQ reduction pathways are required for short-term cold-resistance under Gpx4 dysfunction in hamster cells. (A) Assessment of knock-out efficiency of ferroptosis-suppressors in HapT1 cells. Immunoblots of each protein in non-treated HapT1 (WT) or HapT1 cell populations, in which lentivirus vectors expressing SpCas9 and sgRNA (non-targeting control; sgNT, or sgRNA targeting each of three genes) were infected. (B-D) The proportion of dead cells during cold culture (4°C) in the each HapT1 cell population infected with lentivirus vectors expressing SpCas9 and sgNT or sgRNA for FSP1 (B) or Dhodh (C) or Gch1 (D) in the presence or absence of 2µM RSL3 (One-way ANOVA with the Tukey’s multiple comparison test, p < 0.05). (E) Biopterin synthesis and related pathways. (F) (upper panel) Experimental time course and (lower panel) the proportion of dead cells during cold culture in WT or Gch1 KO HapT cells, which were infected with lentivirus vectors expressing SpCas9 and sgNT or sgRNA for Gch1, in the presence of 6µM ML210 with or without 100µM BH2 and 4µM methotrexate (MTX). (G) Immunoblot of parental HT1080 and HT1080 cell populations in which FSP1 or Gch1 were exogenously overexpressed at 37°C by lentivirus vector infection. (H) The proportion of dead cells during cold culture in parental HT1080 and HT1080 cell populations in which FSP1 or Gch1 were exogenously overexpressed (One-way ANOVA with the Dunnett’s multiple comparison test, ***p < 0.001).

Among the three genes, we decided to focus on the Gch1 and biopterin synthesis pathways because Gch1 KO cells exhibited the highest cell death rate on day2 after treated with Gpx4 inhibitors (Fig. 5d, S2c). Both Pts and Spr encode enzymes that catalyze reactions to form BH4 downstream of Gch1 in the biopterin synthesis pathway (Fig. 5e). A single KO of Pts or Spr in HapT1 phenocopied that of Gch1 (Fig. S2d-f). Next, we investigated the role of Dhfr, which reduces oxidative biopterin BH2 to BH4, in the cold resistance of HapT1. The Dhfr inhibitor methotrexate (MTX) did not affect the survival of HapT1 cells in the absence of Gpx4 inhibitors (Fig. S2g). Dual inhibition of Dhfr and Gpx4 in HapT1 cells did not increase cold-induced cell death compared to Gpx4 inhibition alone (Fig. 5f). These results suggest that the enzymatic activity of Dhfr at 4°C, if any, does not contribute to the cold resistance of HapT1. On the other hand, supplementation of BH2 1hour before cold treatment completely inhibited the cold-induced cell death of both WT and Gch1 KO cells in the presence of ML210, but this effect was abolished by treating the cells with MTX 2hour before cold treatment (Fig. 5f). This suggests that the reduction of exogenous BH2 to BH4 by Dhfr at 37°C before cold treatment is necessary to prevent cold-induced cell death induced by Gpx4 inhibition.

We further addressed whether overexpression of FSP1, Dhodh, or Gch1 could render cold resistance to non-hibernator cells, as well as Gpx4. We infected HT1080 cells with a lentivirus vector that introduced each of these genes derived from hamsters to obtain bulk HT1080 cell lines that overexpressed them (Fig. 5g). Overexpression of Dhodh caused growth defects and did not allow the cell line to be obtained. Overexpression of Gch1 greatly suppressed cold-induced ferroptosis for 6 days, whereas overexpression of FSP1 did not (Fig. 5h).

Finally, we examined whether these ferroptosis suppressors confer cold resistance to mouse primary hepatocytes (Fig. 6a, 6b). Mouse hepatocytes overexpressing maGpx4 via AAV infection showed a significantly higher survival rate than the control cells on day1 (Fig. 6c). Moreover, the administration of BH2 and/or αT with Gpx4 overexpression further augmented the cold resistance of mouse hepatocytes (Fig. 6c). Taken together, BH4 synthesis and CoQ reduction pathways contribute to the cold resistance of hibernator hamster cells in parallel with Gpx4, and overexpression of Gch1 or biopterin supplementation can suppress cold-induced cell death in non-hibernators such as human and mouse.

**Fig. 6.**
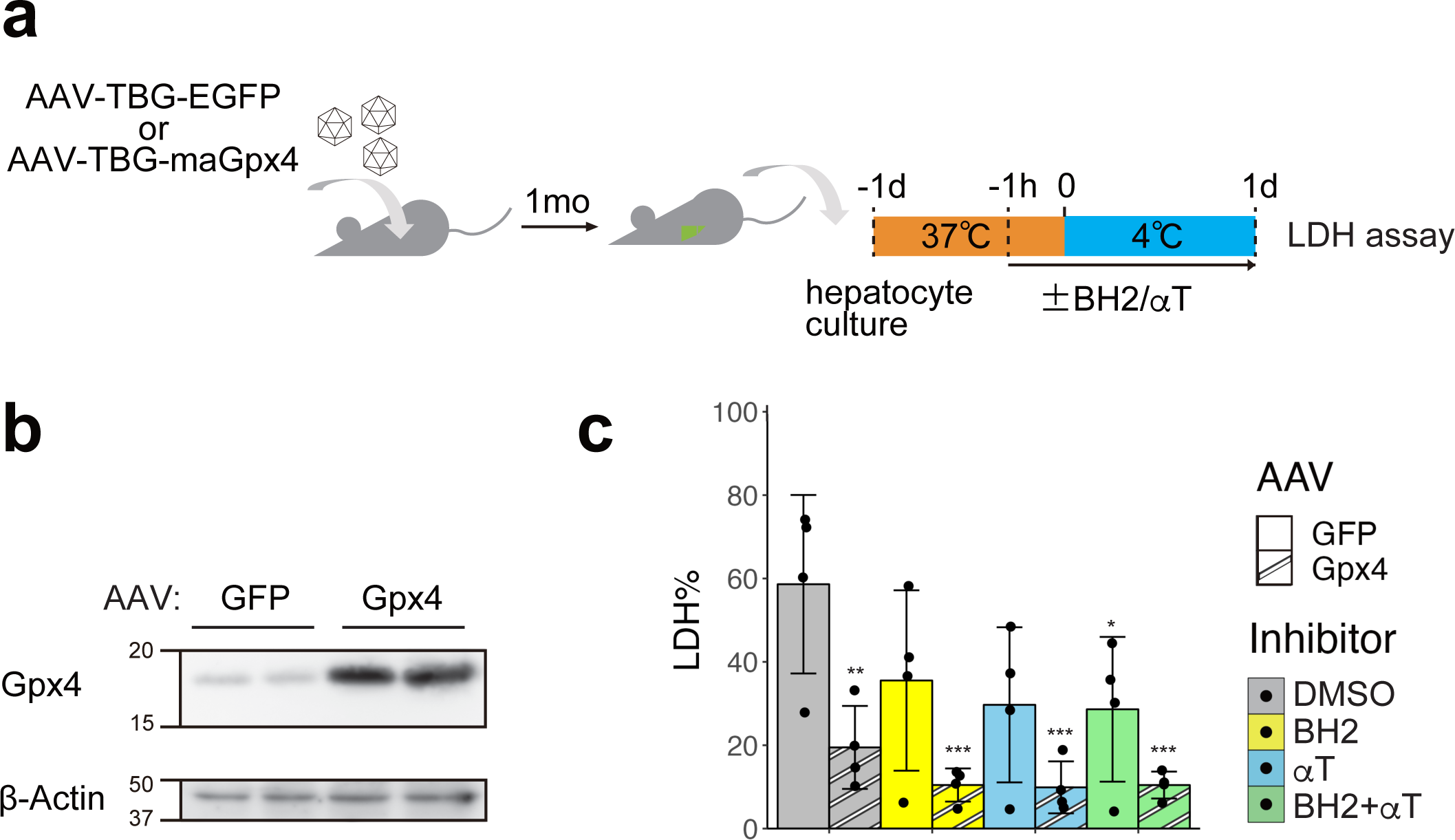
Gpx4 OE and RTAs additively augment cold resistance of mouse primary hepatocytes. (A) Schematic illustration of experimental time course. (B) Overexpression of Gpx4 protein was confirmed by immunoblot of hepatocytes infected with the AAVvectors expressing GFP or maGpx4. (C) The proportion of dead cells in hepatocytes infected with the AAV vectors expressing GFP or maGpx4 after 24hr cold culture with DMSO or 200µM BH2 and/or 10µM αTocopherol (αT) (One-way ANOVA with the Dunnett’s multiple comparison test compared to GFP-DMSO, *p < 0.05, **p < 0.01, ***p < 0.001).

## Discussion

For improvement of therapeutic hypothermia and better organ preservation under cold temperatures, mammalian hibernation is an ideal nature model of cold resistance. How mammalian hibernators prevent cold-induced cell death, whereas non-hibernators, including humans and mice, are vulnerable to it, is a long-standing and fundamental question. To address this, we used an unbiased screening approach and identified Gpx4 as a gene that confers cold resistance to cold-vulnerable human cell line and mouse primary hepatocytes. This is the first report to empirically demonstrate that Gpx4 is sufficient for rendering cold resistance to non-hibernator human and mouse cells and is necessary for the intrinsic cold resistance of mammalian hibernator Syrian hamster cells. These findings are plausible considering that cold-induced cell death resembles ferroptosis and that Gpx4 plays pivotal roles in preventing ferroptosis in a wide variety of mammalian tissues and cells, such as the liver (34), kidney (35) and spinal motor neurons (36).

Loss of Gpx4 function in hamster cell lines by genetic or pharmacological methods abolished cold resistance for a long term (5days) that is approximately the same length as a deep torpor bout of hibernating hamsters. In addition, genetic depletion of Gch1, Dhodh, and FSP1 from HapT1 increased cell death induced by short-term cold treatment under Gpx4 inhibition. These results indicate that biopterin synthesis and plasma membrane/mitochondrial CoQ reduction pathways suppress lipid peroxide accumulation under cold conditions in parallel with Gpx4. Classically, BH4 is well appreciated as a redox cofactor of several enzymes such as nitric oxide synthase (NOS) and tyrosine hydroxylase (Th), but it has recently been known to affect a wider redox state in cells independently of its role as a cofactor (37, 38). It has been reported that knockdown of Gch1 in an endothelial cell line led to increased ROS production from mitochondria, largely through a NOS-independent effect (39). Deficiency of the BH4 synthesis pathway sensitizes human cell lines to ferroptosis inducers, and an in vitro reconstitution assay using liposomes revealed that BH4 directly or indirectly reduces lipid peroxide radicals via αT (22). Additionally, in hepatic carcinoma cells, BH4 suppresses the stress-sensitive Ask1-p38 MAPK signaling pathway (40), which is activated by cold-induced lipid peroxidation in human cell lines (10). Collectively, we propose that BH4 contributes to the cold resistance of hibernator cells not as a cofactor of enzymes but as an RTA reducing lipid peroxide radicals in the membrane. Dhodh is reported to suppress ferroptosis because its inhibition by genetic disruption or administration of BQR at high concentrations (∼1mM) sensitize cells to Gpx4 inhibitor (24). However, questions regarding the interpretation of these results were raised by the finding that the ferroptosis-promoting effect of high-dose BQR largely relies on FSP1 inhibition but not on Dhodh inhibition and that the effect of Dhodh KO on sensitization to RSL3 is much smaller than that of FSP1 KO (41). In our case, a single KO of either Dhodh or FSP1 in HapT1 attenuated short-term cold resistance under conditions of Gpx4 dysfunction (Fig. 5C-D), indicating that each of these CoQ-reducing enzymes functions as a suppressor of cold-induced cell death, at least in this hamster cell line.

We propose that the enzymatic activity of Gpx4 at cold temperatures such as 4°C is required to prevent cold-induced cell death based on the following observations: (i) the duration of pharmacological inhibition of Gpx4 in cold is correlated with the proportion of dead cells (Fig. 3G), (ii) inhibition of Gpx4 before cold treatment did not enhance the cell death rate under cold conditions (Fig. 3G), and (iii) Gpx4-KO HapT1 cells accumulated more lipid peroxide than WT cells during cold treatment (Fig. 3D, 3E). This is in striking contrast to Dhfr, whose activity is required for cell survival by reducing BH2 to BH4 before, but not during, cold treatment when Gpx4 does not function (Fig. 5F). Given that enzyme activity generally declines with a decrease in temperature, according to the Arrhenius equation, the enzymatic activity of Gpx4 may be more robust under cold temperatures; that is, it has a lower Q10 value than other general enzymes.

It is unclear why the human Gpx4 protein can prevent cold-induced cell death similar to the hamster protein when its abundance is high (Fig. 4). Among the eight mammalian Gpx family proteins, Gpx4 has the oldest origin (42). Animal Gpx4 was derived from a single ancestral protein common to fungi, bacteria, and plants and gave rise to all the other Gpx proteins in the animal kingdom (42). Therefore, one possible answer to the above question is that the ability of Gpx4 to protect cells from cold-induced ferroptosis may be preserved in poikilothermic organisms for a long history of life and is inherited to present mammals. In addition, transgenic mice overexpressing mitochondrial Gpx4 in the heart maintained cardiac functionality better after ischemia reperfusion (I/R) than WT animals (43). Accumulating evidence suggests that I/R causes massive ROS production and leads to ferroptosis in thrombosis, such as in stroke and myocardial infarction (44, 45). Several hibernators exhibit resistance to I/R stress (46, 47), though it has not been elucidated whether cold resistance and I/R resistance share common molecular mechanisms. Hibernators’ resistance against I/R stress can be recapitulated by oxygen-glucose deprivation in vitro model (48), indicating cell intrinsic mechanisms to prevent I/R-induced ferroptosis. One study reported that under oxygen-deprived conditions, primary renal cells from hamsters expressed Gpx4 at a higher level than those from mice (49). However, we did not observe any increase in the amount of Gpx4 protein in HapT1 cells after cold exposure (data not shown), and in the livers from animals in hypothermic deep torpor compared with others in euthermic periodic arousal or in the non-hibernation period (Fig. S1c). Therefore, our data suggest that regulation of Gpx4 protein levels is not a mechanism to avoid cold-induced cell death in the hamster cell line and in organs, at least in liver, during hibernation.

It is still an open question why non-hibernator cells are much more vulnerable to cold stress than hibernator cells, even though the expression levels of Gpx4 protein are comparable, as is the case with HT1080 and HapT1 cells (Fig. 2a, Fig. S1a). As evident from the BODIPY time-course analysis, under cold temperatures, lipid peroxidation occurred far more rapidly in HT1080 than in WT HapT1 or Gpx4-KO HapT1. Thus, HT1080 produced a large amount of lipid peroxides, far beyond the capacity of endogenous Gpx4 for reduction (Fig. 3d, 3e). As such, in hibernator cells, Gpx4-independent ferroptosis-suppressing pathways, including BH4-or CoQH_2_-mediated pathways, may decrease the production rate of lipid peroxide under cold temperatures to the level at which endogenous Gpx4 can extinguish it. Such a larger capacity of these Gpx4-independent pathways to prevent lipid peroxidation would provide higher cold resistance to hibernators compared to non-hibernators. Another possible explanation is that hibernator cells may have a superior ability to sustain glutathione levels continuously under cold conditions, as demonstrated in our and previous studies (9), thereby providing reducing power to Gpx4 and other enzymes that utilize glutathione to combat ROS under cold conditions and to prevent lipid peroxidation (Fig. 1f). Our metabolomic analysis revealed that, unlike most other amino acids, cysteine was maintained and tended to increase under prolonged cold and nutrient-deprived conditions. Methionine, a source of cysteine, was rapidly depleted. These data implicate activation of the protein-derived cystine-cysteine supply pathway, possibly by autophagy.

In normal cellular activity, mitochondria consume most of the O_2_ consumed in the cells, and several lines of evidence suggest that mitochondria are one of important candidates for producing ferroptosis-triggering ROS in cold conditions. At cold temperatures, the mitochondria of cells derived from non-hibernators exhibit abnormally high or low membrane potential, whereas those derived from hibernators sustain membrane potential (7, 8). Genome-wide loss-of-function screening using cold-vulnerable human cancer cell lines revealed that disruption of MICU1, which positively regulates the transport of Ca^2+^ ions into the mitochondrial matrix, prevents cold-induced cell death (32). In addition, depletion of intracellular Ca^2+^ by BAPTA-AM in human cancer cells strongly suppressed cold-induced cell death, and this effect was not observed in erastin-induced ferroptosis at 37°C (32). These data suggest that unlike drug-induced ferroptosis at 37°C, cold-induced cell death could be caused by intracellular Ca^2+^ abnormalities leading to mitochondrial dysfunction and ROS production, which eventually oxidizes lipids in the plasma membrane. Considering that Gpx4 and other ferroptosis-suppressors exist in non-hibernators that nevertheless fail to exert cold resistance, it is an interesting topic for future studies to address how hibernators maintain Ca^2+^ homeostasis at cold temperatures.

Collectively, we clarified that hibernator cells can sustain glutathione synthesis pathway under cold conditions and prevent cold-induced ferroptosis through the concerted action of Gpx4-dependent lipid peroxide reduction, biopterin synthesis, and CoQ reduction pathways. We also demonstrated that overexpression of Gpx4 or increasing intracellular BH4 levels was sufficient to render cold resistance in non-hibernator cells. A limitation of this study is that the contribution of these ferroptosis suppressors to in vivo cold resistance during hibernation was not tested. Future studies will address this point through genetic manipulation of these pathways in hamsters using CRISPR/Cas9 or RNAi technologies, which are now feasible for this animal (25, 50). Other approaches, such as genome-wide loss-of-function screening in hibernator cells, may also help to elucidate crucial components for cold resistance of hibernators other than known ferroptosis-related genes. An additional crucial aspect to consider is the possible oncogenic consequences of altering ferroptosis-associated genes, as they are potential targets of anti-cancer treatments. Hence, it will be necessary to investigate approaches to mitigate any such effects in the context of clinical applications. In conclusion, this study is the first step in addressing the causal relationship of genes and cold resistance of hibernators, one of the long-lasting mysteries of hibernation, thereby providing novel clues for the improvement of clinical applications, such as therapeutic hypothermia and cold preservation for organ transplantation.

## Acknowledgements

We thank Hirotaka Imai (Kitasato University), Jun Suzuki (Kyoto University), Hiroshi Yamaguchi (Nagoya University), Genshiro Sunagawa (Riken BDR), Masayuki Miura (University of Tokyo), and all members of the Yamaguchi Laboratory for their helpful discussions regarding this study.

## Author Contributions

MS, YS and YY contributed to the design and implementation of the research, MS, NM, YS, YM, RM, AY, RO, JY, KS, SE and DA to the performance of experiments and the analysis of the results, MS, YS and YY to the writing of the manuscript.

