## Supplemental figure and legends for "Identification of genes supporting cold resistance of mammalian cells: lessons from a hibernator"

### Supplemental figure legends

**Fig. S1 Expression levels of Gpx4 in HT1080, HapT1 and liver tissues of mice and hamsters**

(a) Immunoblot of Gpx4 and β-Actin proteins in HT1080 and HapT1.

(b) Immunoblot of Gpx4 and β-Actin proteins in mouse and hamster livers (N = 4).

(c) Immunoblot of Gpx4 and β-Actin proteins in livers of hamsters in the states of non-hibernation (room temperature, RT), periodic arousal (PA) and deep torpor (DT) (N = 3 or 4). Signals were quantified and represented as box plot in the right panel.

**Fig. S2 Biopterin and CoQ reduction pathways are required for short-term cold-resistance under Gpx4 dysfunction in hamster cells, related to Fig. 5**

(a-c) The proportion of dead cells during cold culture (4℃) culture in the each HapT1 cell population infected with lentivirus expressing SpCas9 and non-targeting sgRNA (sgNT) or sgRNA for FSP1 (a) or Dhodh (b) or Gch1 (c) in the presence or absence of 6μM ML210 (One-way ANOVA with the Tukey's multiple comparison test, $p < 0.05$).

(d) Assessment of knock-out efficiency of Spr in HapT1 cells. Immunoblots of Spr and β -Actin proteins in HapT1 cell populations infected with lentivirus vectors expressing SpCas9 and sgNT or sgRNA targeting Spr.

(e) Assessment of knock-out efficiency of Pts in HapT1 cells. Quantitative RT-PCR of Spr and β-Actin mRNA in HapT1 cell populations infected with lentivirus vectors expressing SpCas9 and sgNT or sgRNA targeting Pts. Note that Pts mRNA was decreased in the cells introduced with sgPts presumably through nonsense mediated RNA decay pathway.

(f) The proportion of dead cells after two-day cold culture in HapT1 infected with lentivirus expressing SpCas9 and sgNT or sgRNA for Spr or Pts and treated with 2μM RSL3 or 6μM ML210 determined by LDH assay (One-way ANOVA with the Dunnett's multiple comparison test, $^{**}p < 0.01$, $^{***}p < 0.001$).

(g) The proportion of dead cells during cold culture in HapT1 infected with lentivirus expressing SpCas9 and sgNT or sgRNA for Gch1 with or without 100μM BH2 and 4μM methotrexate (MTX) determined by LDH assay (N = 3 wells).

Fig.S1

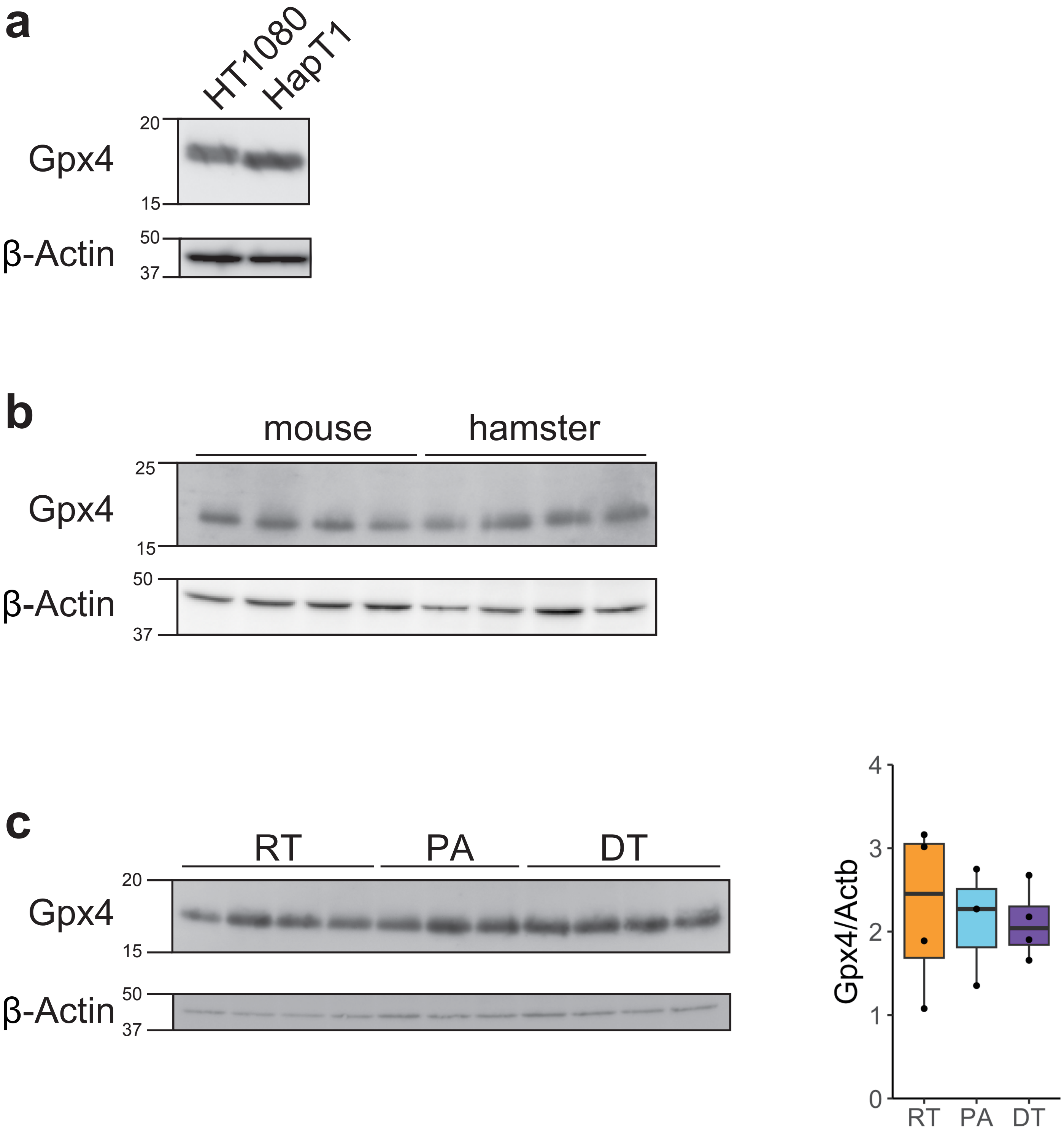
a
HT1080
HapT1
Gpx4
β-Actin
20
15
50
37
b
mouse
hamster
Gpx4
β-Actin
25
15
50
37
c
RT
PA
DT
Gpx4
β-Actin
20
15
50
37
Gpx4/Actb
0
1
2
3
4
RT
PA
DT

Fig. S2

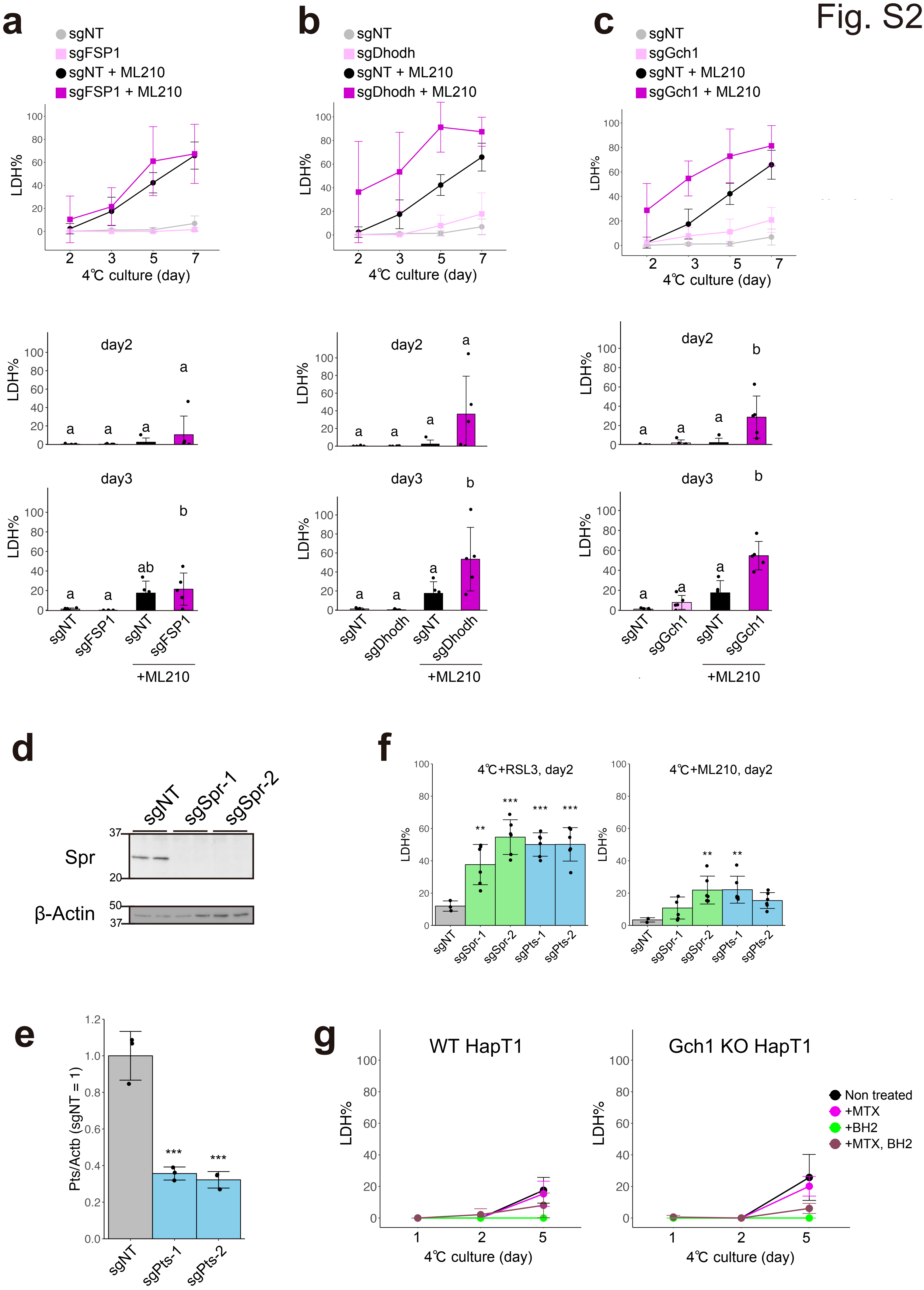

a
sgNT
sgFSP1
sgNT + ML210
sgFSP1 + ML210
LDH%
4℃ culture (day)
day2
day3
sgNT
sgFSP1
+ML210
b
sgDhodh
sgDhodh + ML210
c
sgGch1
sgGch1 + ML210
d
sgNT
sgSpr-1
sgSpr-2
Spr
β-Actin
e
Pts/Actb (sgNT = 1)
sgPts-1
sgPts-2
f
4℃+RSL3, day2
4℃+ML210, day2
g
WT HapT1
Gch1 KO HapT1
Non treated
+MTX
+BH2
+MTX, BH2
